# Stress-Induced Expression is Enriched for Evolutionarily Young Genes in Diverse Budding Yeasts

**DOI:** 10.1101/660274

**Authors:** Tyler W. Doughty, Iván Domenzain, Aaron Millan-Oropeza, Noemi Montini, Philip A. de Groot, Rui Pereira, Jens Nielsen, Céline Henry, Jean-Marc G. Daran, Verena Siewers, John P. Morrissey

**Affiliations:** Department of Biology and Biological Engineering, Chalmers University of Technology, SE-41296, Gothenburg, Sweden; Novo Nordisk Foundation Center for Biosustainability, Chalmers University of Technology, SE-41296, Gothenburg, Sweden; Plateforme d’Analyse Protéomique Paris Sud-Ouest (PAPPSO), INRA, MICALIS Institute, Université Paris-Saclay, 78350 Jouy-en-Josas, France; School of Microbiology, University College Cork, T12YN60, Ireland, Centre for Synthetic Biology & Biotechnology, APC Microbiome Ireland, and Environmental Research Institute, Cork, Ireland; Department of Biotechnology, Delft University of Technology, Van der Maasweg 9, 2629 HZ Delft, The Netherlands

## Abstract

The Saccharomycotina subphylum (budding yeasts) spans more than 400 million years of evolution and includes species that thrive in many of Earth’s harsh environments. Characterizing species that grow in harsh conditions could enable the design of more robust yeast strains for biotechnology. However, tolerance to stressful conditions is a multifactorial response, which is difficult to understand since many of the genes involved are as yet uncharacterized. In this work, three divergent yeast species were grown under multiple stressful conditions to identify stress-induced genes. For each condition, duplicated and non-conserved genes were significantly enriched for stress responsiveness compared to single-copy conserved genes. To understand this further, we developed a sorting method that considers evolutionary origin and duplication timing to assign an evolutionary age to each gene. Subsequent analysis of the sets of genes that changed expression revealed a relationship between stress-induced genes and the youngest gene set, regardless of the species or stress in question. These young genes are rarely essential for growth and evolve rapidly, which may facilitate their functionalization for stress tolerance and may explain their stress-induced expression. These findings show that systems-level analyses that consider gene age can expedite the identification of stress tolerance genes.

## Introduction

Yeasts in the Saccharomycotina subphylum, “budding yeasts”, have proven to be useful platforms for the production of ethanol, flavors, nutritional supplements, biopharmaceuticals, as well as other valuable chemicals^1,2,3^. At present, industrial production using budding yeasts as a platform is dominated by the extensively characterized species *Saccharomyces cerevisiae. S. cerevisiae* exhibits common budding yeast phenotypes (e.g. efficient growth on simple sugars) as well as a less common adaptation amongst budding yeasts, high ethanol tolerance^4^. Together, these traits enable cost-effective production of 100 billion liters of ethanol annually using *S. cerevisiae* as a platform^1^. Other budding yeasts have adaptations that make them well-suited for production of specific biomolecules, something that is possible due to the improved strain engineering capacity following the emergence of CRISPR/Cas9^5,6^. Examples are *Yarrowia lipolytica,* which evolved to tolerate hydrophobic environments and can produce high-yields of fatty acids^7,8^, and *Kluyveromyces marxianus,* whose thermotolerance is a beneficial feature for industrial processes^6,9^. Despite progress in sequencing genomes and phenotypic characterization of these and many other yeast species, the genes that underpin adaptation to cope with harsh conditions remain enigmatic.

For the species above, adaptation to natural environments enables robustness in industrial biotechnology processes. Understanding an exceptional stress tolerances from specific budding yeast species, might therefore enable the engineering of pan-robust industrial strains, thereby reducing process costs and increasing yields^10,11^. Although studies that sought to characterize stress tolerances in *S. cerevisiae* have elucidated mechanisms that influence robustness^10,12,13^, engineering more robust *S. cerevisiae* strains without physiological trade-offs remains challenging^9^. One complication is that stress exposure often results in hundreds of significant transcriptional changes^13,14^, most of which do not correlate with single gene deletion changes in robustness^11^. These results suggest that several genes from different gene families may contribute additively to robustness and/or that stress genes may exist as duplicates, as is the case for antifreeze protein genes in artic yeasts^15^. Thus, researchers have employed systems biology to characterize the transcriptome and/or proteome-wide stress-induced changes^13,14,16–18^. These approaches have identified biological processes that exhibit altered expression in response to stress exposure, which build upon and relate to extensive previous research into gene functions (e.g. GO term enrichment analysis). These associations are possible due to extensive annotations of *S. cerevisiae* genes that result from decades of experimental analyses^19^. For most other yeast species, the majority of gene functional information is acquired second hand via homology search tools. This paradigm results in a large portion of genes of unknown function, which is especially large for species that are phylogenetically distant from extensively characterized species like *S. cerevisiae*^20^. These uncharacterized genes are difficult to integrate into omics analyses like GO term enrichment, as they do not have a known function or localization. Because of this, gene functional analysis of poorly characterized species is restricted to conserved genes, which may not be the only genes that influence stress-tolerance phenotypes. Currently, hundreds of whole genome sequences are available from diverse budding yeasts^21^, including several species that are known to exhibit extreme stress tolerances^22^, but many of the causative genes that enable yeast stress tolerances remain elusive.

Here, we analyzed stress conditions to assess gene expression changes after stress adaptation in three diverse budding yeast species, one of which is well characterized (*S. cerevisiae*), and two that are less-well-characterized (*K. marxianus* and *Y. lipolytica*). The goal of this analysis was to identify common systems-level trends that are shared between each species stress responses. This analysis discovered that each organism displayed a consistent response at the level of gene expression that was characterized by the enrichment of stress responsive genes amongst certain categories: namely, to genes of unknown function and to recently (in evolutionary time) duplicated and taxonomically restricted genes (young genes). The findings of this work suggest an evolutionary mechanism that is biased for stress tolerance functionalization and periodic expression of young genes. We propose that the gene sorting method herein provides a path forward for more rapid identification of stress response genes in environmentally robust yeast, thereby accelerating understanding of niche adaption in budding yeasts.

## Results

### Stress Adaptation Responsive Genes are Enriched for Duplicated and Non-Conserved Genes

In this work, *S. cerevisiae, K. marxianus,* and *Y. lipolytica* were exposed to stress conditions that are present in natural environments, such as those caused by seasonal temperature variation and growth on sugar-rich or acidic substrates^22^. These stress responses are industrially-relevant, as they are caused by feedstocks (high osmotic pressure and low pH) or process conditions (elevated temperatures) during industrial fermentations^11^. Characterizing stress responses in these species is valuable due to their phylogenetic diversity, which spans much of the Saccharomycotina subphylum^21^. To minimize noise caused by variable growth rate^23^, experiments were carried out in steady-state chemostats at a fixed growth rate under standard or stress conditions. This experimental setup allows strains to acclimate to, and grow in the presence of sub-lethal stress before sampling and analysis, which is referred to as stress adaptation herein. Transcriptomic changes that occurred in response these stress conditions were identified via differential expression analysis (Figure 1A).

**Figure 1).**
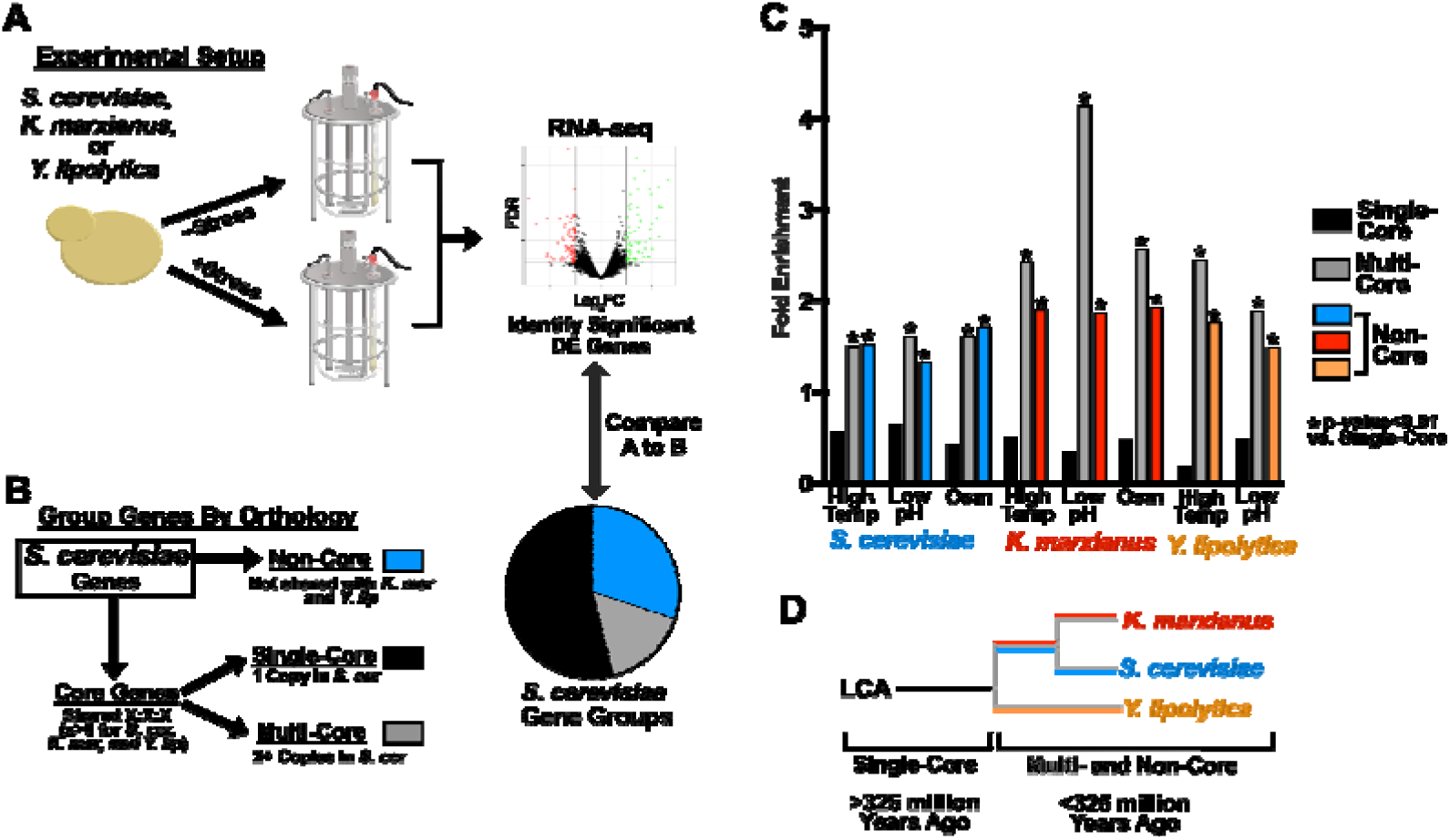
Stress Adaptation Responsive Genes are Enriched for Duplicated and Non-Conserved Genes. A. *S. cerevisiae, K. marxianus,* and *Y. lipolytica* were cultivated in chemostats at a growth rate of 0.1/h in standard conditions, or in the presence of stress (elevated temperature, low pH, or KCl). RNA-seq was performed followed by differential expression analysis. B. The protein coding genes of *S. cerevisiae, K. marxianus*, and *Y. lipolytica* were probed for orthology using OrthoFinder. *S. cerevisiae* gene grouping is shown as an example. Orthologous genes (core genes) were identified and grouped by their copy number in *S. cerevisiae* with multi-copy orthologs in gray and single-copy orthologs in black. Genes that were not matched to orthologs were designated as non-core genes (blue). C. Differentially expressed (log_2_FC>1, FDR<0.01) mRNAs were normalized to the total number of detected genes inside of their respective group resulting in a percentage DE for each gene group, which was normalized to the DE % of the dataset to obtain fold enrichment. D. A diagram of expected gene origin for the species in this figure. Single Core orthologs are predicted to originate from a Last Common Ancestor >325 million years ago. Multi- and Non-Core Genes are predicted to have duplicated, arisen *de novo,* or via another evolutionary event less than 325 million years ago.

In order to understand the function of stress responsive genes, biological process annotations were acquired from Ensembl (*S. cerevisiae*) or identified using BLAST2GO^20^ for (*K. marxianus* and *Y. lipolytica*). BLAST2GO annotated gene functions to otherwise unknown genes based on homology to an experimentally characterized gene. This process failed to annotate 20% and 40% of the mRNAs measured by RNAseq in this study for *K. marxianus* and *Y. lipolytica* respectively (Supplemental 1A). The lower frequency of gene annotation for *Y. lipolytica* was expected, since this species is not closely related to extensively characterized yeasts^21^. Comparison of gene annotations and differential gene expression showed a higher percentage of genes of unknown function that were stress responsive than would be expected. For example, 38% of all protein-coding genes measured in this study for *Y. lipolytica* lacked a functional annotation, while 50% of stress responsive genes were genes of unknown function (Supplemental Figure 1B).

This high proportion of stress-responsive genes of unknown function suggested that the most broadly conserved genes, which often have functional annotations, might be under-represented amongst the stress responses. To assess this, orthologous proteins shared between the three yeast species were inferred using OrthoFinder, which enables proteome-wide matching based on amino-acid sequence and chain length similarity in order to predict proteins that descend from a common ancestor^24^. To assess the fidelity of ortholog predictions, protein complexes and enzymatic processes that are expected to be conserved amongst budding yeasts were searched for amongst orthology inference results^25^. This analysis found that orthology inference identified the majority of the expected complex members and enzymes as orthologs (Supplemental Figure 2B), which supports the high fidelity of OrthoFinder predictions that was observed previously^24^. The results of the orthology inference analysis were used to divide each protein into one of three classes, single-core orthologous, multi-core orthologous, and non-orthologous. These proteins were matched to their corresponding genes for comparison to RNAseq differential expression. Gene sorting examples are shown in Supplemental Figure 2A and the entire list of genes can be found in Supplemental Sheet I.

**Figure 2).**
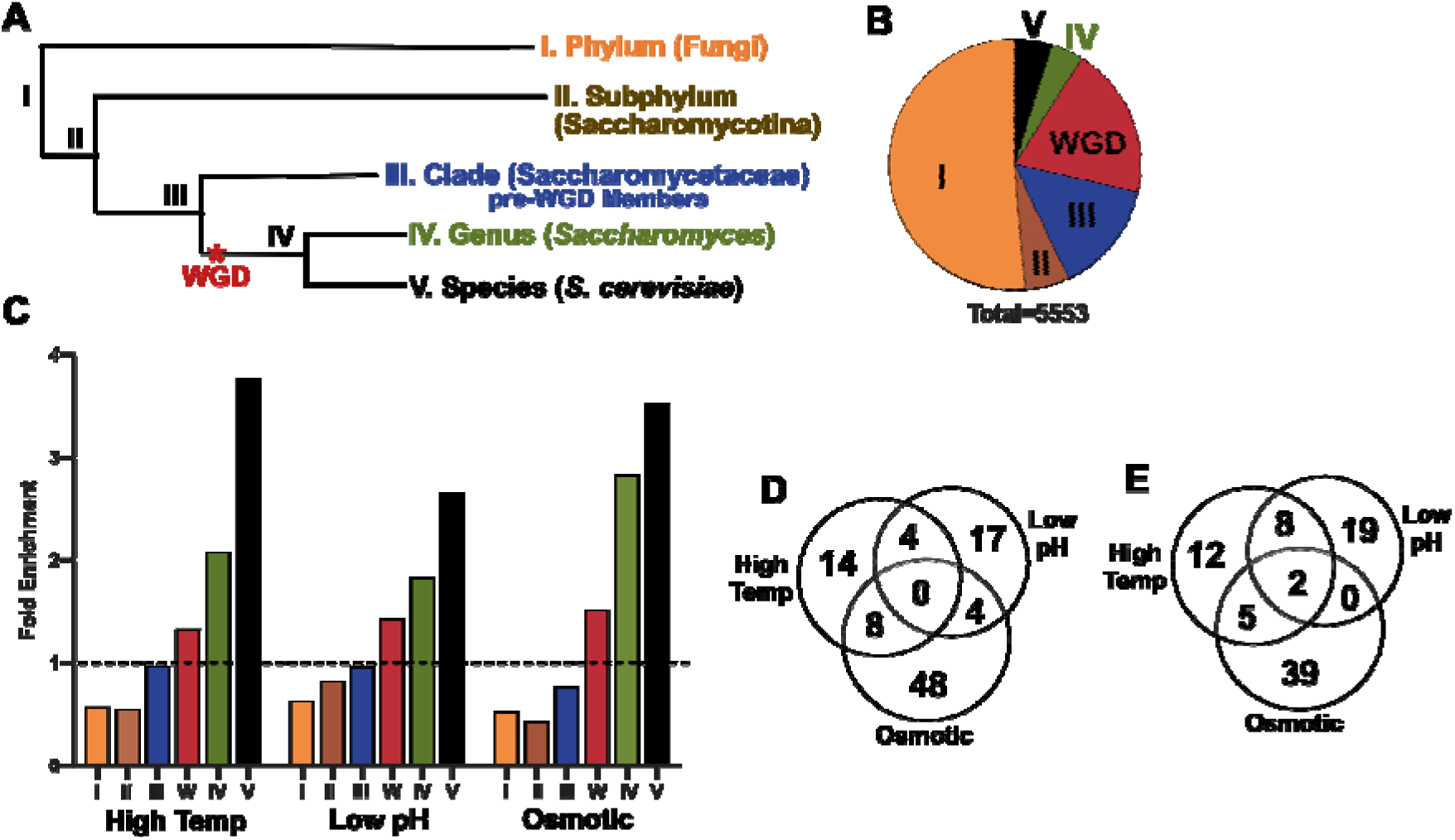
Stress Adaptation Responsive Genes in *S. cerevisiae* are Enriched for Young Genes. **A** A simplified phylogenetic tree for *S. cerevisiae* showing speciation events and the Whole Genome Duplication (red *). **B** The measured transcripts from this study were grouped based on gene age via orthology inferrence (described in Supplemental Figure 4 and Supplemental Methods). **C** Differentially expressed genes for *S. cerevisiae* were parsed by their grouping shown in b, then normalized to the group size and the total measured DE % (dashed line). Transcripts in groups IV and V were assessed for shared downregulated genes (**D**) or upregulated genes (**E**).

The results of orthology inference for *S. cerevisiae* are shown in Figure 1B as an example. Each measured protein-coding gene from *S. cerevisiae* was identified as either 1) present as a single-copy gene with an ortholog in *K. marxianus* and *Y. lipolytica* (black Single-Core), 2) present as a duplicated gene with an ortholog in *K. marxianus* and *Y. lipolytica* (gray Multi-Core), or 3) lacking an ortholog in *K. marxianus* or *Y. lipolytica* (color Non-Core). The resulting groups were compared to the observed differentially expressed genes, which showed that multi-core and non-core genes were significantly enriched amongst DE genes in each stress condition tested (Figure 1C). The same gene sorting regime shows that *K. marxianus* and *Y. lipolytica* exhibited similar DE gene enrichment for the multi-core and non-core gene groups (Figure 1C and Supplemental Figure 3A). Similar results were found amongst proteomics measurements for some stress conditions, but this analysis was hindered by low detection of non-core proteins (Supplemental Figure 3C).

**Figure 3).**
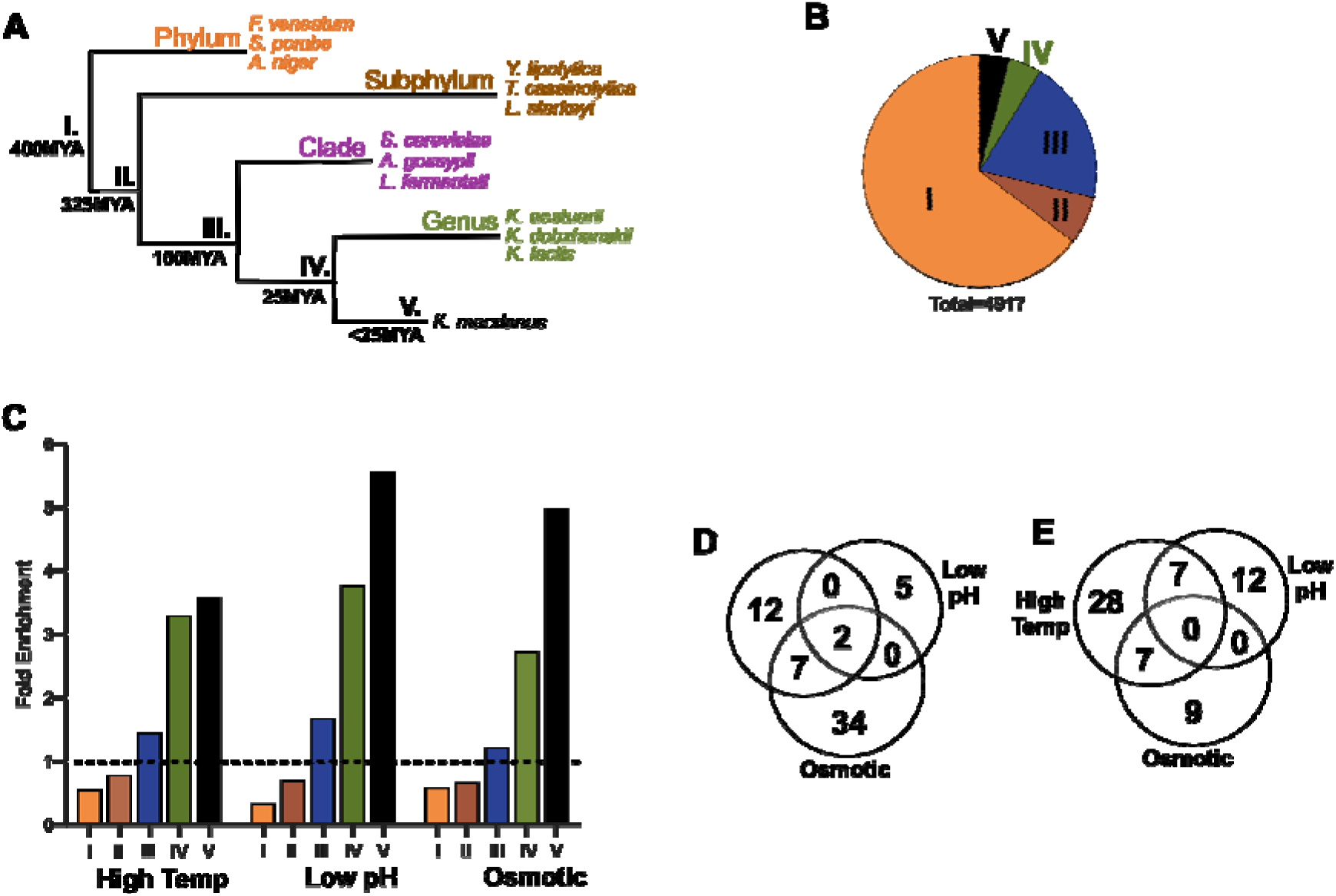
Stress Adaptation Responsive Genes in *K. marxianus* are Enriched for Young Genes. **A** A simplified phylogenetic tree for *K. marxianus* showing speciation events and organisms used in orthology queries. **B** Detected transcripts from this study were grouped based on ortholog presence in the groups shown (described in supplemental Figure 4). **C** Differentially expressed genes for *K. marxianus* were parsed by their grouping shown in A and B, then normalized to the group size and the total measured DE % (dashed line). Transcripts in groups IV and V, were assessed for shared downregulated genes (D) or upregulated genes (E).

The phenomenon depicted in Figure 1C shows that single-core genes, which are predicted to have descended from a last common ancestor between the three yeast species (approximately 325 million years ago^21^) were under-represented amongst stress responsive genes for each stress and each organism. In contrast, genes that have duplicated or emerged in more recent evolutionary time were enriched amongst stress responsive genes. These observations suggest that evolutionary events may predict differential expression amongst these diverse yeast species (Figure 1D).

### Stress Adaptation Responsive Genes in *S. cerevisiae* are Enriched for Young Genes

The results in Figure 1 suggested a relationship between the genes that exhibit differential expression in response to stress and evolutionary events, like *de novo* gene emergence and gene duplication. Further characterization of this relationship could aid in understanding stress gene evolution and could help to predict novel genes that enable stress tolerance. Thus, we sought to test this relationship more stringently by dividing the protein-coding genes of *S. cerevisiae* into more precise groups that collectively represent a broad swath of eukaryotic evolution. The resulting groups are referred to as gene age groups, which were determined by ortholog presence at shared copy number in common ancestors that date from over 400 million years ago to 20 million years ago^21^. A similar approach, phylostratigraphy, divides genes into groups based on homology and has been used to infer gene origination events to identify periods in evolution that correlate with adaptive events^26^. However, the results in Figure 1C indicated that an analysis paradigm that considers both gene origin timing (like phylostratigraphy) and gene duplication timing could provide new insights into stress responsive gene expression.

Gene grouping based on gene age was assessed using OrthoFinder^24^ and is described in detail in the Supplemental Methods. Briefly, all *S. cerevisiae* genes were divided into three initial subsets; 1) fixed duplicates from the whole-genome duplication (WGD)^27^, 2) genes that are present as single-copy genes, and 3) duplicate genes that arose outside of the whole-genome duplication (non-WGD) (Figure S4A). Ortholog inference was used to sort each of the 4,351 single-copy genes into a single bin based on the most distant ancestor with an orthologous gene using the hierarchal approach shown in Figure S4C. The multi-copy non-WGD gene groups were sorted by the presence of orthologous genes with the same copy number in a bottom-up approach in order to trace the relative timing of gene duplication events (Supplemental Figure 4D). Finally, genes that were duplicated during the whole-genome duplication were grouped together. This sorting method matched each protein coding gene from *S. cerevisiae* to a single group that reflects the timing of the emergence (single-copy genes) or timing of duplication (multi-copy genes) of each gene, which we refer to as gene age. A complete list of genes measured in this study and their relationships is found in Supplemental Table II.

**Figure 4).**
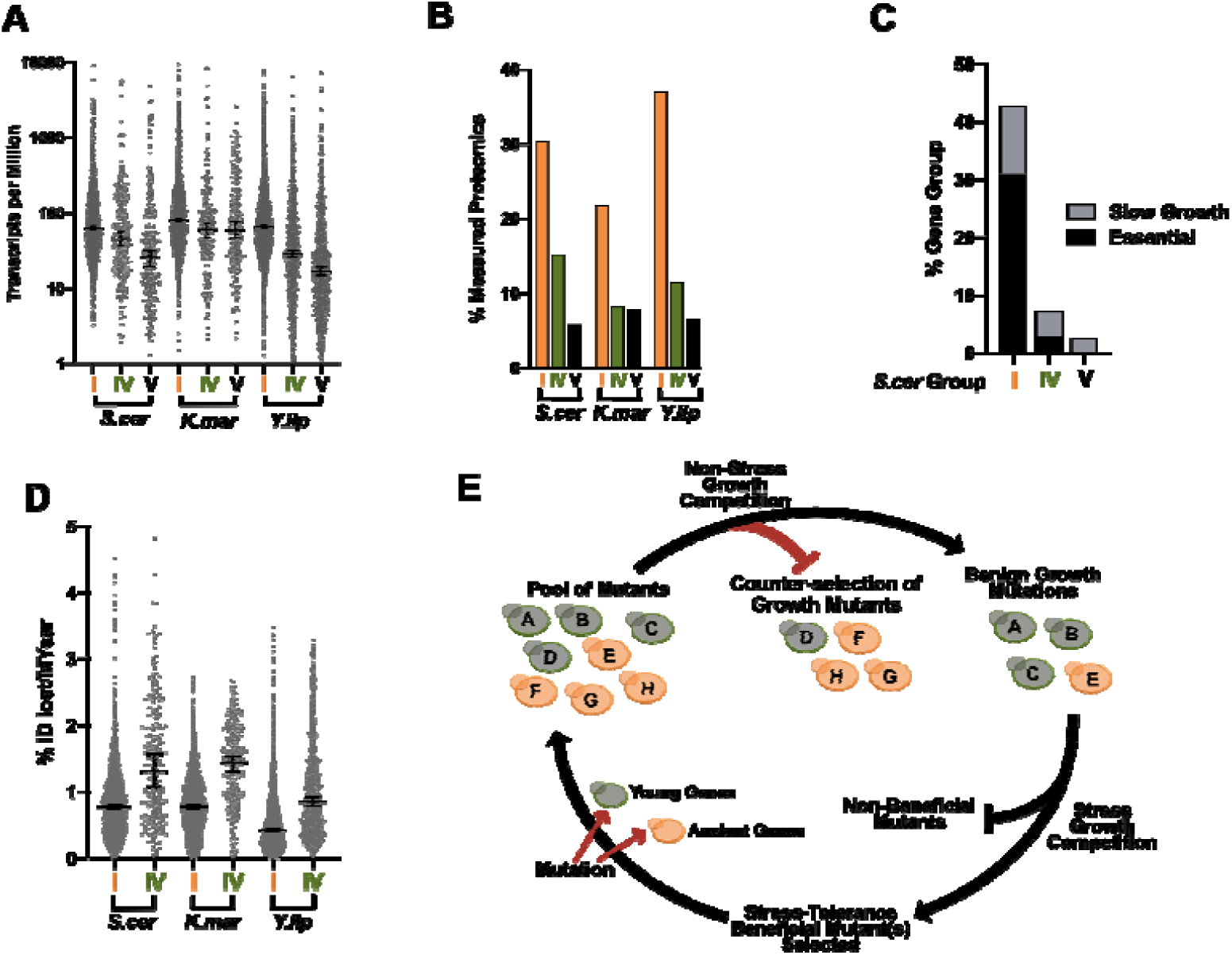
Young Genes are Less Expressed, Often Non-Essential, and Adapt More Rapidly than Ancient Genes. **A.** Transcripts measured in standard growth conditions were normalized to the read depth and gene length to generate Transcripts per Million (TPM). Error bars at the 95% confidence interval of the median. B. The percentage of mRNAs measured in this study compared to proteins measured via mass spectrometry by quantifying eXtracted Ion Chromatograms. Experimental samples from standard conditions were used for this analysis. C. The percentage of essential genes (black) and non-essential genes associated with slow growth (gray) is shown for *S. cerevisiae* ancient genes (I) and young genes (IV and V). Essential and slow growth ORFs were obtained from Giaever 2002^20^. D. The percentage of amino acid identity changes for each protein in comparison to its closest homolog from a member of the same genus. Results were adjusted to the percent amino acid change per million years (% ID lost/MYear) using the estimated divergence time between pairs of organisms^13^. The median and 95% confidence interval is shown for each group is denoted by a black line. Queries were performed between *S. cerevisiae/S. eubayanus, K. marxianus/K. lactis*, or *Y. lipolytica/Y. bubula*. E. A model for evolution to intermittent stress. Random mutations occur naturally amongst genes in each age group (red arrows) followed by non-stress selection for benign mutants (red blocked arrow). Mutants that do not influence growth are selected upon stress exposure for fitness benefits.

The gene groupings in Figure 2B were compared to the stress RNAseq data to determine the percentage of significantly differentially expressed genes in each age group. This analysis found a stepwise increase in differentially expressed gene enrichment in progressively younger gene groups in *S. cerevisiae*. Genes that were found to be conserved to filamentous fungi (ancient genes from group I) were 4.2 to 6.6-fold less likely to be differentially expressed after stress adaptation compared to *S. cerevisiae-*specific genes (group V) (Figure 2C). Similar trends were observed when considering only upregulated or downregulated genes, however, upregulated genes showed a more pronounced bias towards young genes with 6.6 to 16.8 fold enrichment between group I and group V genes (Supplemental Figure 5). Analysis of the expression pattern of young genes (those in groups IV and V) showed that few genes exhibited significantly changed expression in response to all stresses (Figures 2D and 2E).

The findings in Figure 2 were further tested by analyzing additional stress adaptation experiments for *S. cerevisiae* exposed to ethanol in a previous study^28^ or anaerobic stress (this study) (Supplemental Figure 6). In both cases, young genes were enriched, and ancient genes were depleted amongst differentially expressed genes in response to stress adaptation. A similar enrichment for young genes was observed amongst varying amounts of ethanol stress, despite difference in the number of total significant gene expression changes (Supplemental Figure 6D). Together, these observations suggest that gene age is a viable predictor of differential expression amongst many types and levels of stress in *S. cerevisiae*.

### Stress Adaptation Responsive Genes in *K. marxianus* and *Y. lipolytica* are Enriched for Young Genes

The findings in Figure 2 showed an inverse correlation between gene age and stress differential expression in *S. cerevisiae*. If these findings were shared amongst other yeast species, they might imply an underlying evolutionary mechanism that can predict the genes that are more likely to be involved in stress adaptation. To test for a relationship between differential expression and gene age, we stratified the protein-coding genes of *K. marxianus* and *Y. lipolytica* using the same sorting concept described above for *S. cerevisiae* (Supplemental Figure 4). The only modification to these sorting approaches was the elimination of the whole-genome duplication group, as neither of these species has undergone a recent whole-genome duplication^29,30^.

Analysis of *K. marxianus* and *Y. lipolytica* gene groups in relation to each stress condition showed similar patterns to *S. cerevisiae*, with ancient genes exhibiting under-representation for significant differential expression compared to young gene groups (Figure 3 and Supplemental Figure 7). Also, as with *S.* cerevisiae, there were few young differentially expressed genes that responded to all stresses, suggesting that these expression changes were often condition specific (Figure 3D and 3E). These biases towards young genes might explain the low observed overlap between significant expression changes amongst 1:1:1 orthologs shared between the three budding yeasts when exposed to the same type of stress (Supplemental Figure 8). Together, these findings showed that in all three yeasts studied, young genes were enriched for long-term stress-responsiveness, or adaptation, compared to ancient genes. Further, since the species chosen for this analysis span much of the diversity of the budding yeast subphylum^21^, these results may be indicative of a shared stress adaptation mechanism, rather than a shared response of specific genes, amongst budding yeasts.

### Young Genes are Less Expressed, Shorter, Less Likely to be Essential, and Accumulate Sequences Changes More Rapidly than Ancient Genes

To understand the functions associated with the gene groupings produced in this study, we assessed biological process enrichment amongst the ancient and young gene sets in *S. cerevisiae*, where ample functional information is available. This analysis showed ancient genes associated with fundamental biological processes including primary metabolism, tRNA aminoacylation, and DNA strand elongation, and 94% of these genes were annotated with at least one biological process GO term. Conversely, young genes (groups IV and V) were associated with more specialized functions like maltose transport, vitamin biosynthesis, and aldehyde metabolism, with many young genes lacking any biological process annotations in *S. cerevisiae* (40%). *K. marxianus* and *Y. lipolytica* also exhibited high percentages of young genes that were not associated with a biological process (41% and 69% respectively) (Supplemental Figure 9B). The fundamental nature of ancient gene functional associations was reflected by their high likelihood to be essential or required for optimal growth compared to young genes. Conversely, the more specialized functions of young genes were reflected by the 16-fold decrease in likelihood of growth impairment upon deletion compared to ancient genes (Figure 4C)^31^. Analysis of cellular component enrichment showed that young proteins (groups IV and V) were significantly enriched for localization to the plasma membrane, cell wall, and vacuole, which was distinct from ancient proteins (group I) enrichment for nuclear, cytoplasmic, and mitochondrial localization (Supplemental Table III).

Further characterization of young protein-coding genes found that they exhibited lower median gene expression and were less frequently detected via mass spectrometry in non-stress samples compared to ancient genes (Figures 4A and 4B). Previous works have shown that low expression and non-essentiality correlate with increased adaptation rates^32,33^, suggesting that young genes could adapt more rapidly compared to ancient genes. To test this, amino acid sequence identity was compared between homologous proteins from members of the same genus using BLAST+. Analysis of each protein sequence from groups I and IV allowed sequence identity changes to be compared over the same span of evolutionary time to assess adaptation rates. This analysis was adjusted to reflect the estimated evolutionary time elapsed^21^ between each pair of species and showed that the average frequency of amino acid identity changes was higher for young protein groups compared to ancient protein groups (Figure 4D).

## Discussion

Budding yeasts are attractive for industrial production of biomolecules, since they grow rapidly, utilize inexpensive substrates, and are readily engineered to produce heterologous gene products^1,2,3^. However, stresses that result from feedstock composition, toxic products, and fluctuating reaction temperatures can lower the cost-effectiveness of industrial processes by diminishing productivity and yields^11^. Previous works have phenotypically characterized yeasts exhibiting stress tolerant phenotypes^22^, and whole genome sequencing data are available, but the genes that have evolved in these yeasts to enable survival and growth under unfavorable, stress-inducing conditions remain unclear. We now identify an association between stress-induced gene expression and gene age. We show that younger genes, namely, those that have are restricted to a genus or species, or have duplicated in recent evolutionary time, are more likely to respond to different types of long-term stress, such as those that we imposed in continuous (chemostat) cultivation in this report. These stress-responsive genes can also be considered adaptation or niche-specialization genes as they evolved to enable the yeasts carrying them tolerate ongoing harsh conditions.

The findings that adaptation rates and stress gene expression are biased toward young genes for three distantly related yeast species suggests an underlying evolutionary mechanism. The model in Figure 4E suggests that during non-stress periods, ancient and young gene mutations may occur at similar rates, however, ancient genes may be subject to more stringent counter-selection (red blocked arrow) due to their higher expression and influence on growth (Figures 4A and 4C). Conversely, non-synonymous mutations amongst young genes might accumulate more rapidly because these genes are rarely growth-related (Figure 4C and 4D). The resulting increase in sequence space that is sampled by young genes would increase the probability of young mutants to enter stress-growth competition, thus increasing the chances of selecting young gene adaptations to benefit stress tolerance. We suggest that these events occur in a cyclical manner, enabling stress-tolerance functionalization of young genes without diminishing growth potential. This model could also apply to promoter sequences, which would enable specialized genes to adapt dynamic expression patterns in order to save resources during non-stress growth. This mechanism would explain the higher propensity of young genes to change expression in response to stress. The model might also provide an insight as to why improved stress tolerance in some laboratory-evolved strains comes at a cost to growth under standard growth conditions^34,35^. In this case, the relatively short, non-cyclical stresses applied during adaptive laboratory evolution does not allow for the counterselection of growth mutations.

In this work we found that young genes represented 4%, 5%, and 14% of protein-coding genes in *K. marxianus, S. cerevisiae,* and *Y. lipolytica*, respectively, which is in the same range as the 7-19% of genes in *C. elegans, D. melanogaster,* and *H. sapiens* that lack recognizable homologs in other organisms^26,36^. Previous works have linked some young genes to species and genus-specific adaptations, including movement on the surface of fast water in *Rhagovelia* water striders^37^, HIV-1 resistance in owl monkeys^38,39^, and the concurrent evolution of antifreeze proteins in several species^40–42^. Antifreeze protein genes are well-studied examples of young genes that arose via *de novo* gene origin events between 13 and 18 million years ago in codfishes and are present at variable copy number in some species^43^. Concurrently, the psychrophilic yeast *G. antarctica*, has evolved to encode nine antifreeze protein genes whose expression levels are induced by exposure to cold^15,44^. These attributes of antifreeze protein genes are similar to the young genes in this study, which were stress responsive, emerged in recent evolutionary time, and often exist at variable copy number. It seems plausible that the young, stress responsive genes described herein for *K. marxianus* could influence this species growth at higher temperatures (45°C)^9^ than other members of the *Kluyveromyces* genus, like *K. lactis* (37°C)^45^. Further, the acquisition of this thermotolerant phenotype in a short span of evolutionary time would be consistent with the involvement of rapidly adapting young genes.

This study and previous stress tolerance investigations have identified several dozen to several hundred significant gene expression changes after stress exposure in budding yeasts^13,16–18,28^. Despite analysis of stress-responsive genes in several robust species, rational engineering of robust strains remains difficult. The findings of this work suggest that perturbing evolutionarily young stress-responsive genes from stress tolerant species is a pragmatic path forward toward designing robust industrial strains. This suggestion is based on two points; 1) single gene perturbations often fail to reproduce stress-response phenotypes^13^ and 2) many mutations that improve stress tolerance cause trade-off phenotypes^10,34,35^. Establishing more robust industrial production strains may require several gene mutations and/or expression of several exogenous genes, while avoiding growth or physiological perturbations. To accomplish this, new, knowledge-driven approaches are needed to aid the identification of relevant genes that can be manipulated to confer the desired trait without negative consequences on growth. These aims are complicated by incomplete gene function information, which is particularly lacking for many stress tolerant yeast species. In this work, we present a gene sorting method that identifies a class of genes that are likely to be enriched in response to diverse stresses. By leveraging gene age information, it will be possible to focus rational experimental designs on novel stress tolerance genes. Identifying these genes offers biotechnological potential as well as the tools to understand the process of species diversification and niche adaptation in yeast.

## Methods

### Strains and Cultivation Conditions

*Y. lipolytica* (W29), *K. marxianus* (CBS6556), and *S. cerevisiae* (CEN.PK113-7D) were grown in 30mL synthetic media at 30°C for 24 h in shake flasks, followed by inoculation of bioreactors and an initial batch growth phase. After the completion of the batch phase, chemostat cultivation was started with a dilution rate of 0.1/h and a working volume of 500 mL (*S. cerevisiae*) or 1 L (*K. marxianus* and *Y. lipolytica*). Stress conditions were achieved by altering either temperature, pH, or osmotic pressure (KCl) for the duration of the cultivation, specific conditions are listed in Supplemental Figure 8. Standard growth temperature was adjusted to reflect organism specific tolerances. Cultivations for were performed in synthetic medium (SM)^46^ containing 5 g·L^-1^ (NH_4_)_2_SO_4_, 3 g·L^-1^ KH_2_PO_4_, 0.5 g·L^-1^ MgSO_4_·7H_2_O, 7.5 g.L^-1^ glucose, trace elements and vitamins with 1 g L^-1^ pluronic PE6100 to reduce foaming. Sample collection was carried out after at least five volume changes (50 h) in steady state growth conditions. Steady state growth was defined as less than 5% deviation in biomass dry weight.

### Ortholog Prediction with OrthoFinder

For Figure 1, proteome-wide homology matching was executed using OrthoFinder^24^. Proteins were excluded from the core genome (non-core) if orthology search predicted zero orthologous proteins in any of the query species. Proteins were designated single-core if they were encoded by single-copy genes in the species (e.g. *S. cerevisiae HIS1*) or multi-core if they were duplicated in the species (e.g. *S. cerevisiae GAL1* and *GAL3*) (Supplemental Figure 2). Protein groups were matched to their underlying genes for gene expression analyses. This grouping strategy was carried out to sort each species protein-coding genes into a single group. Results of these gene sorting analyses are shown in Supplemental Sheet I. For Figure 2, Figure 3, and Supplemental Figure 7, OrthoFinder was used to identify orthologs between each yeast and a set of eukaryotic organisms. This is shown in Supplemental Figure 4 and is discussed in more detail in the supplemental methods, results of these gene sorting analyses are shown in Supplemental Table II.

### RNAseq Preparation and Mapping

RNA extractions were performed on samples that were mechanically lysed with 0.5mm acid washed beads using an MP-Biomedicals™ FastPrep-24 for three one-minute cycles. Further extraction was performed using an RNeasy® Kit from Qiagen. Libraries were prepared using the TruSeq mRNA Stranded HT kit. Sequencing was carried out using an Illumina NextSeq 500 High Output Kit v2 (75 bases), with a minimum of 8 million paired-end reads per replicate. The Novo Nordisk Foundation Centre for Biosustainability (Technical University of Denmark), performed the RNA sequencing and library preparation. RNAseq datasets can be found using SRA accession PRJNA531619. RNAseq read mapping was performed after analysis in FASTQC, which identified one sample from *K. marxianus* as having overrepresented sequences. This sample was excluded from the analysis herein. Analysis for TPM in Figure 4a was performed using Hisat2^47^ and StringTie^48^. RNAseq mapping for differential expression was mapped with STAR^49^ and reads were assigned with featureCounts^50^. Differential expression results can be found in Supplemental Table 3.

### Mass Spectrometry Measurements for Proteomics

Sample preparation details can be found in supplementary data. Peptides were analyzed in an Orbitrap Fusion™ Lumos™ Tribrid™ mass spectrometer (Thermo Fisher Scientific) in data-dependent acquisition mode using High Collision Dissociation (details in supplementary data). Protein identification was performed using the search engine X!Tandempipeline 3.4.4^51^. Data filtering was set to peptide E-value < 0.05, protein log(E-value) < –2 and to a minimum of two identified peptides per protein. Relative quantification of protein abundance was carried out using two complementary methods, one based on spectral counting and other considering the area of eXtracted Ion Chromatograms (XIC) of a given peptide at specific retention time as described in Millan-Oropeza 2017^52^. MS data are available online on public databases via the PRIDE^53^ repository with the dataset identifier PXD011426 and PROTICdb^51,54^ or via the following URLs:

http://moulon.inra.fr/protic/chassy_kluyveromyces

http://moulon.inra.fr/protic/chassy_yarrowia

http://moulon.inra.fr/protic/chassy_saccharomyces

### Differential Expression Analysis

Differential expression data was generated using limma^55^ and edgeR^56^ R packages, with filtering to remove lowly expressed genes/proteins. In addition, each dataset was filtered to remove genes/proteins for which the relative standard deviation was greater than 1 (RSD>1) for replicates for a given condition and organism. Differential expression was defined by a significance cutoff of absolute log_2_FC>1, False Discovery Rate<0.01 for a stress condition compared to control. All mapped transcript data, protein detection data, custom tools, and analysis scripts can be found at:

https://github.com/SysBioChalmers/OrthOmics

## Supporting information

Supplemental Table III

