## Supplemental Figures and Methods for "Stress-Induced Expression is Enriched for Evolutionarily Young Genes in Diverse Budding Yeasts"

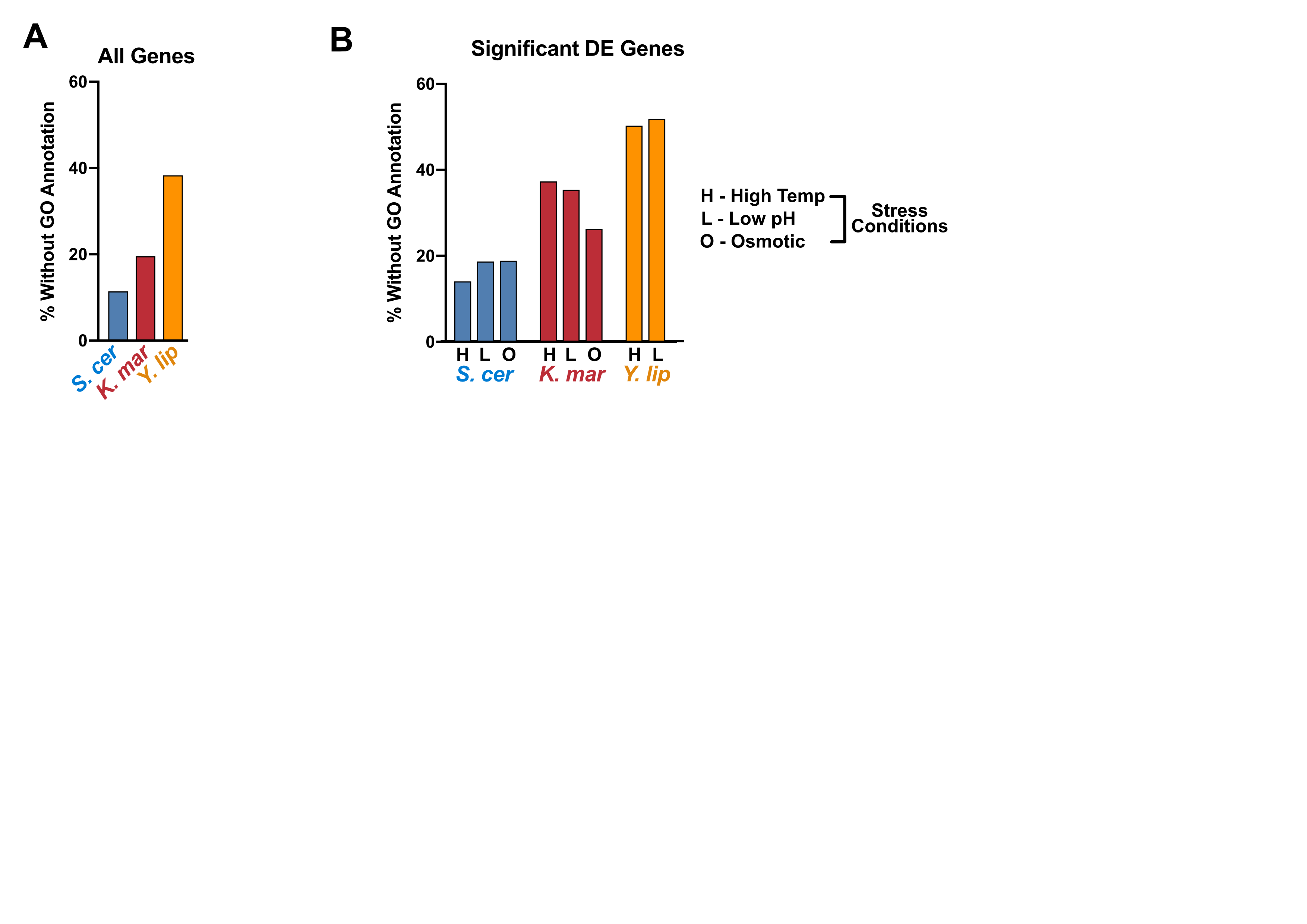

**Supplemental 1) BLAST2GO Annotation of *K. marxianus* and *Y. lipolytica* Leaves Many Stress Responsive Genes without a Functional Annotation.**

A) Ensembl GO terms for *S. cerevisiae* compared to BLAST2GO^1^ annotated GO terms for *K. marxianus* and *Y. lipolytica* showing the % of protein-coding genes detected in this study that lack any biological process annotation. B) Differentially expressed genes in this study compared to the GO annotations used in A.

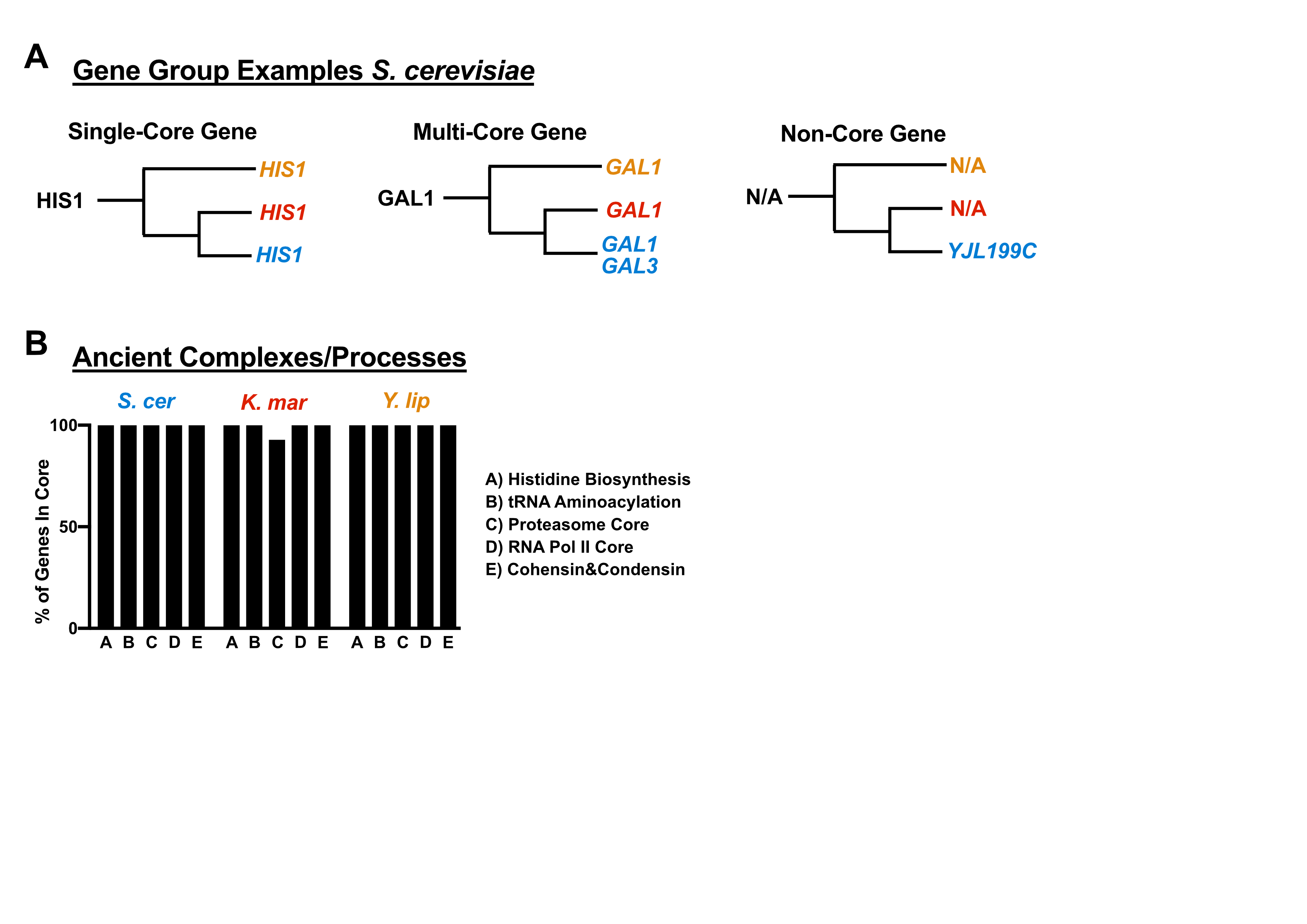

**Supplemental 2) Gene Grouping by Orthology Inference for Gene Groups in Figure 1**

A. Genes from *S. cerevisiae* are shown as examples to explain the gene grouping analysis executed for Figure 1. B. Eukaryotic gene/protein groups that are expected to have evolved before the emergence of the budding yeast subphylum^2^ are shown with the percentage of single-core genes that were identified in Figure 1.

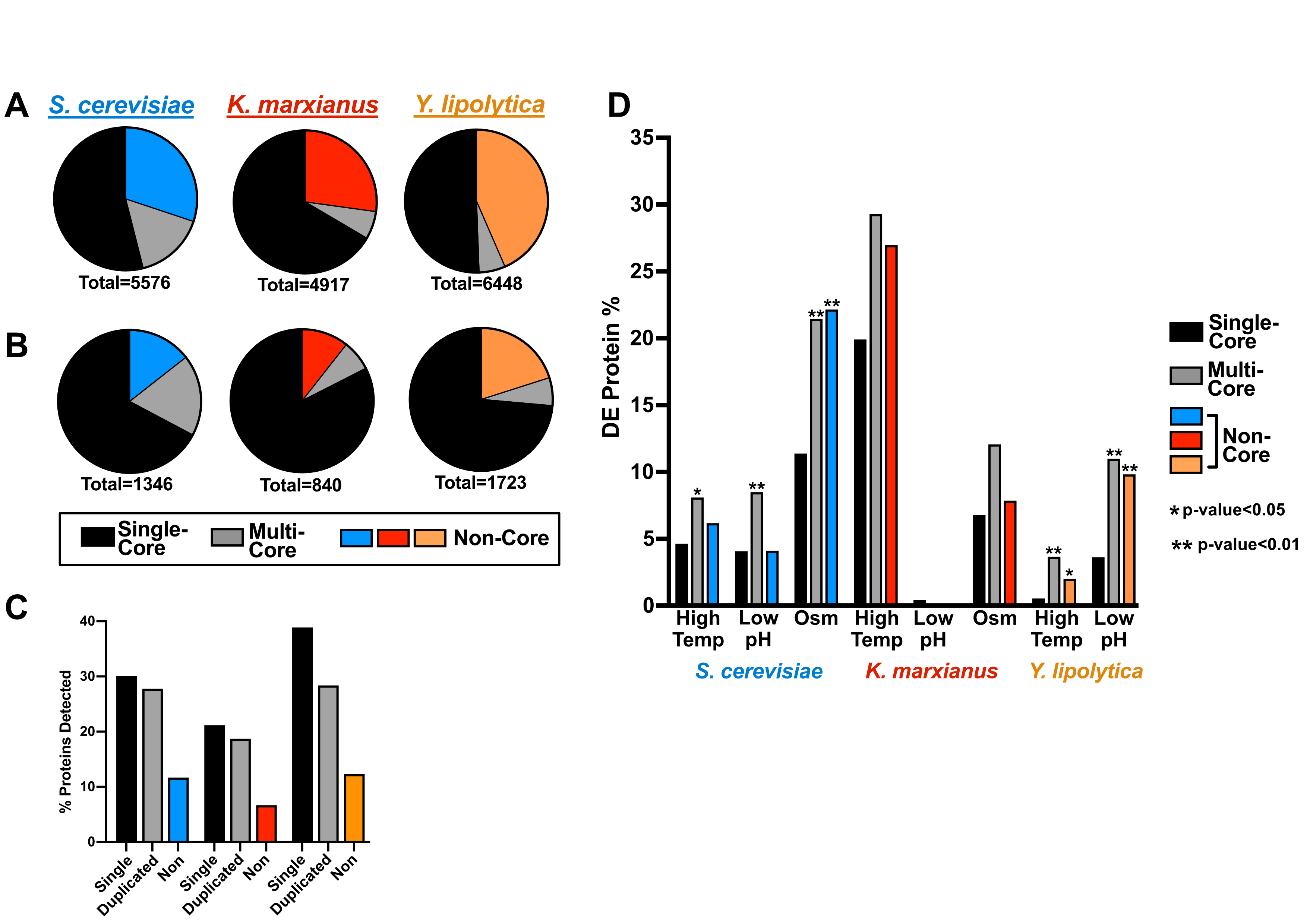

**Supplemental 3) Single-Core Proteins are Depleted for Stress-Responsiveness in Some Conditions, but Analysis is Limited by Low Detection of Non-Core Proteins.**

The transcripts measured via RNAseq (A) and proteins measured via relative proteomics (B) (spectral counts and XIC) were classified into core genes/proteins (black) between the three fungi or non-core genes/proteins (color) using OrthoFinder^3^. C. The number of proteins detected was normalized to the number of mRNAs detected. D. The number of differentially expressed (logFC>1, FDR<0.01) proteins were normalized to the total number of measured proteins inside of their respective group; single copy orthologs (black), multi-copy -orthologs (gray), and non-orthologs (color).

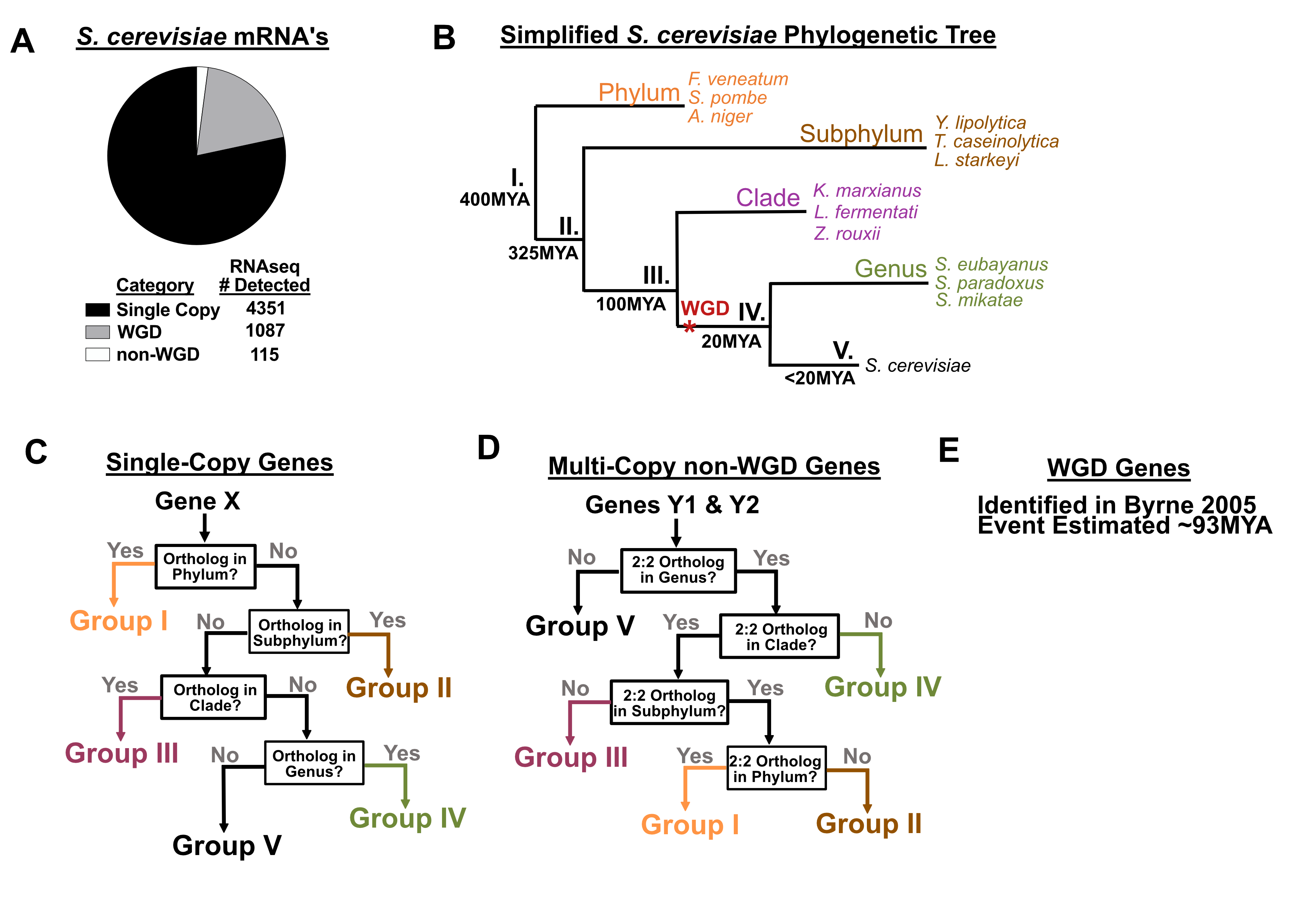

**Supplemental Figure 4) Grouping of *S. cerevisiae* Protein-Coding Genes Using Ortholog Inference Results.**

A The protein-coding genes of *S. cerevisiae* were divided into whole genome duplicates (described in Byrne and Wolfe 2005^4^) and non-WGD genes. Non-WGD genes were scanned for duplications via self-self orthology search, which identified 115 measured genes that had at least one duplicate in *S. cerevisiae*. B. A simplified phylogenetic tree is shown with the names of the species used for the gene sorting queries in C and D. The 4351 single copy genes from A were subjected to ortholog queries against each of the organisms listed. The approximate times of last common ancestors were estimated in previous works^5,6^ (MYA=million years ago). D Non-WGD duplicates gene families were queried for the presence of the same copy number in other related species using the logic shown. All ortholog queries utilized OrthoFinder^3^ and all protein ortholog inferences were applied to the underlying genes for analysis in Figure 3.

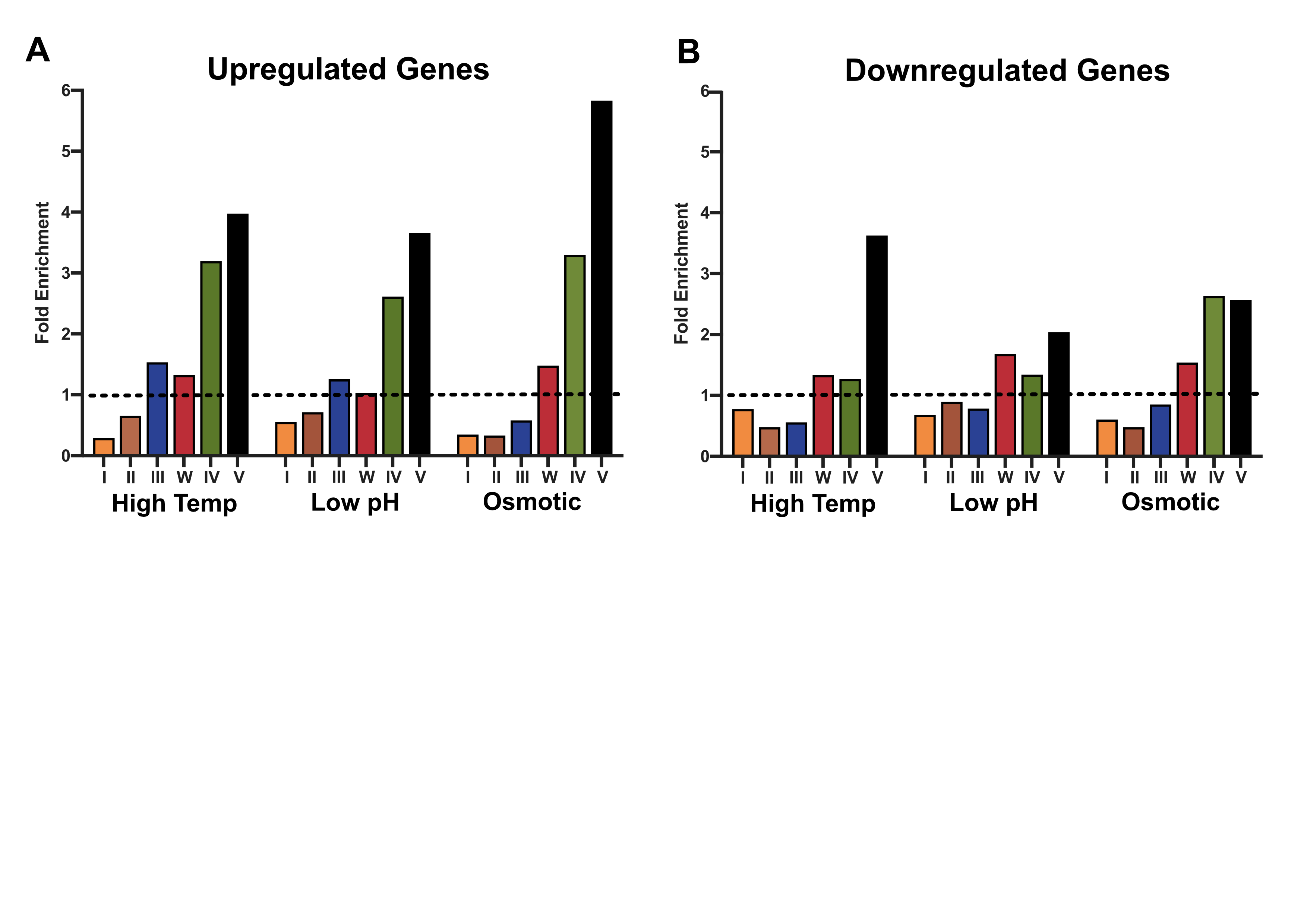

**Supplemental 5) The Stress Adaptation Response in *S. cerevisiae* is Enriched for Both Up and Downregulation of Young Genes.**

A For comparison, the significant gene expression results shown in Figure 3C were divided into upregulated genes (log2FC>1, FDR<0.01) (A) and downregulated genes (log2FC<-1, FDR<0.01) (B).

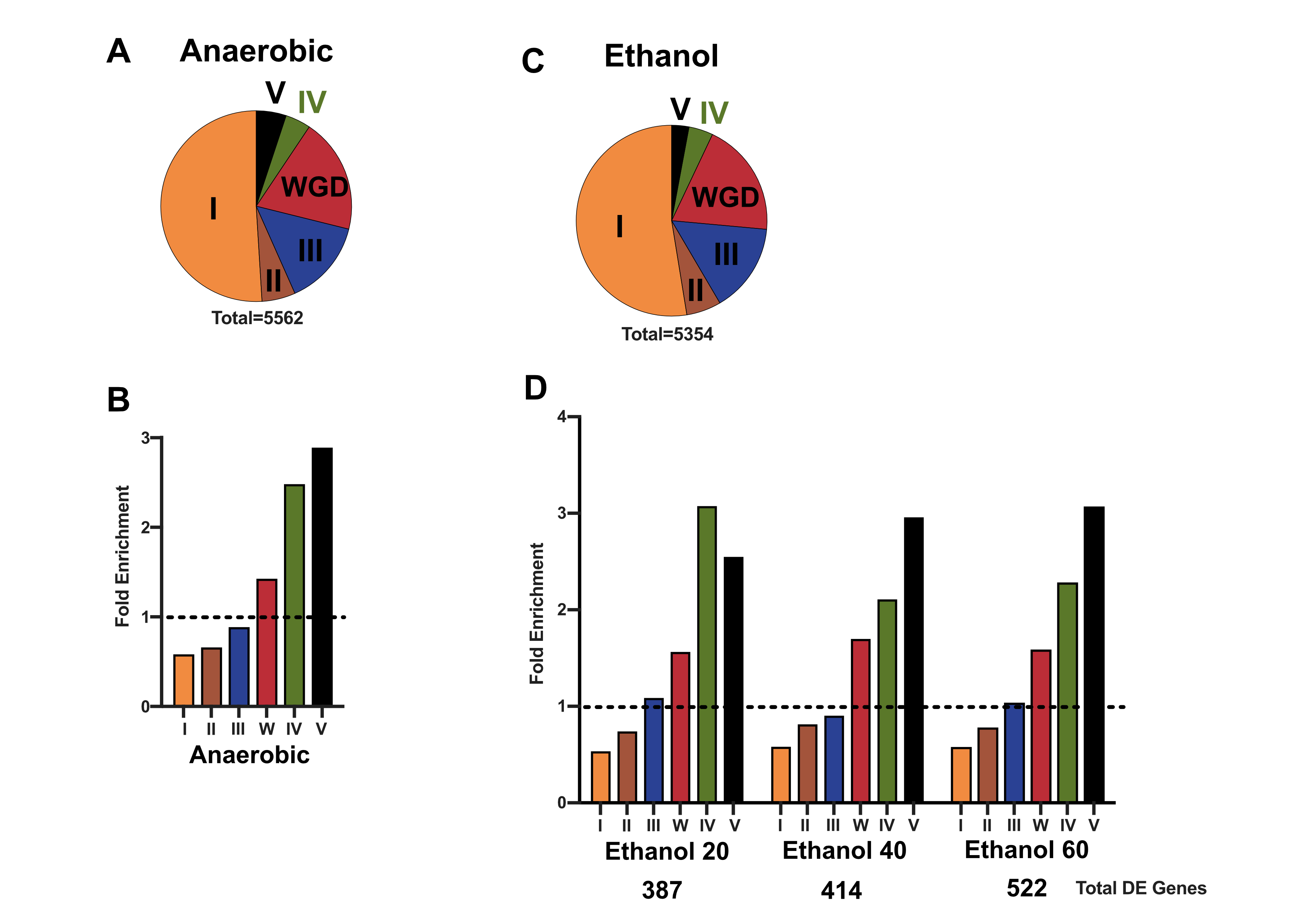

**Supplemental 6) The Stress Adaptation Response in *S. cerevisiae* to Anoxia and Ethanol are Enriched for Young Genes**

Gene groupings were created using the methodology described in Supplemental figure 4 for genes measured in anaerobic stress (A) or ethanol stress (C). Differentially expressed genes for *S. cerevisiae* were parsed by their grouping, then normalized to the group size and the total measured DE % (dashed line) upon anaerobic growth (B) or after exposure to 20, 40, or 60g/L ethanol (D) (dataset from Lahtvee *et al.,* 2017)^7^.

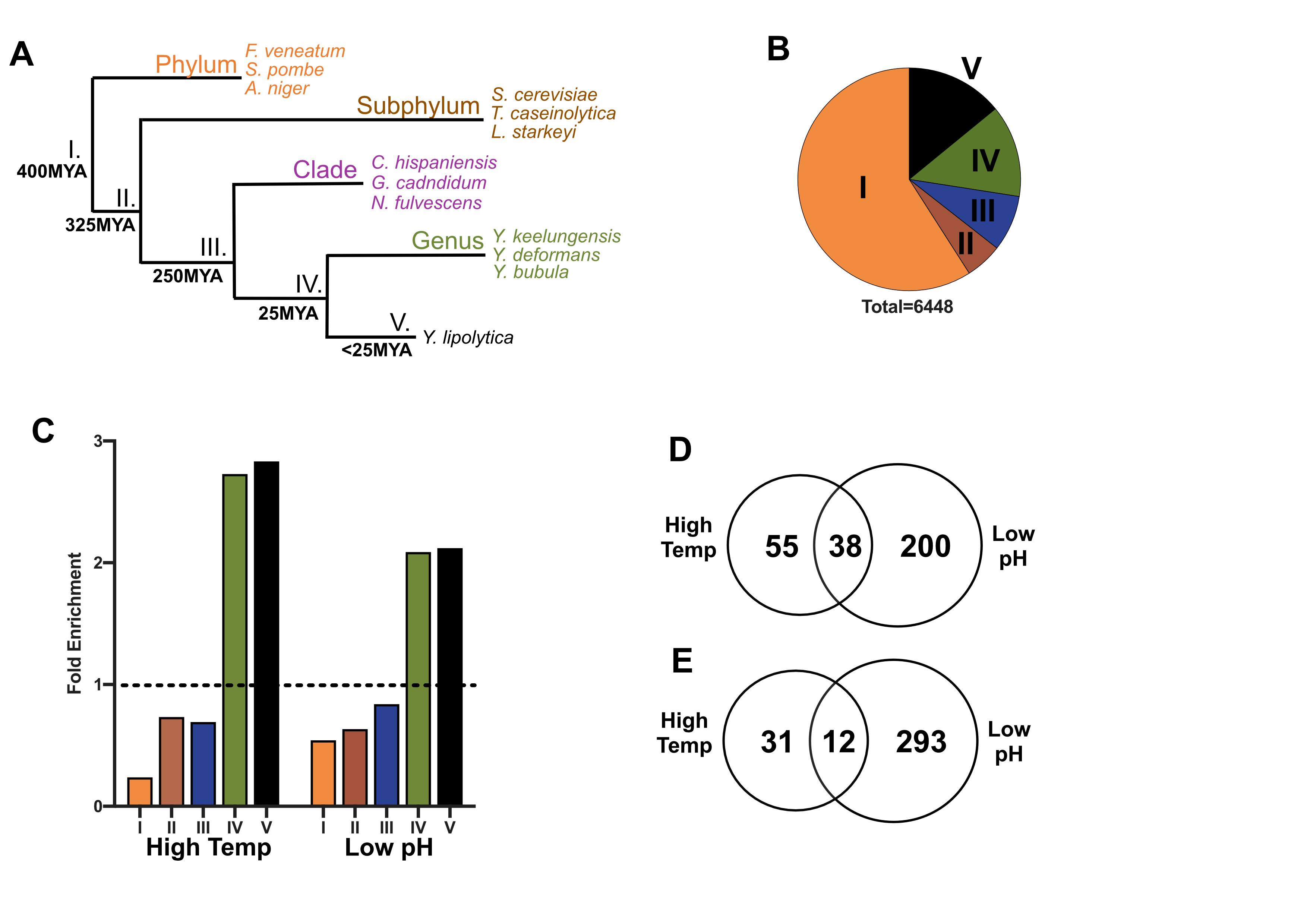

**Supplemental Figure 7) Stress Adaptation Responsive Genes in *Y. lipolytica* are Enriched for Young Genes.**

A A simplified phylogenetic tree for *Y. lipolytica* showing speciation events. B The single-copy transcripts were grouped based on ortholog presence in the groups shown (described in supplemental Figure 4). C Differentially expressed genes for *Y. lipolytica* were parsed by their grouping shown in B, then normalized to the group size and the total measured DE % (dashed line). Transcripts in group IV and V, were assessed for shared downregulated genes (D) or upregulated genes (E).

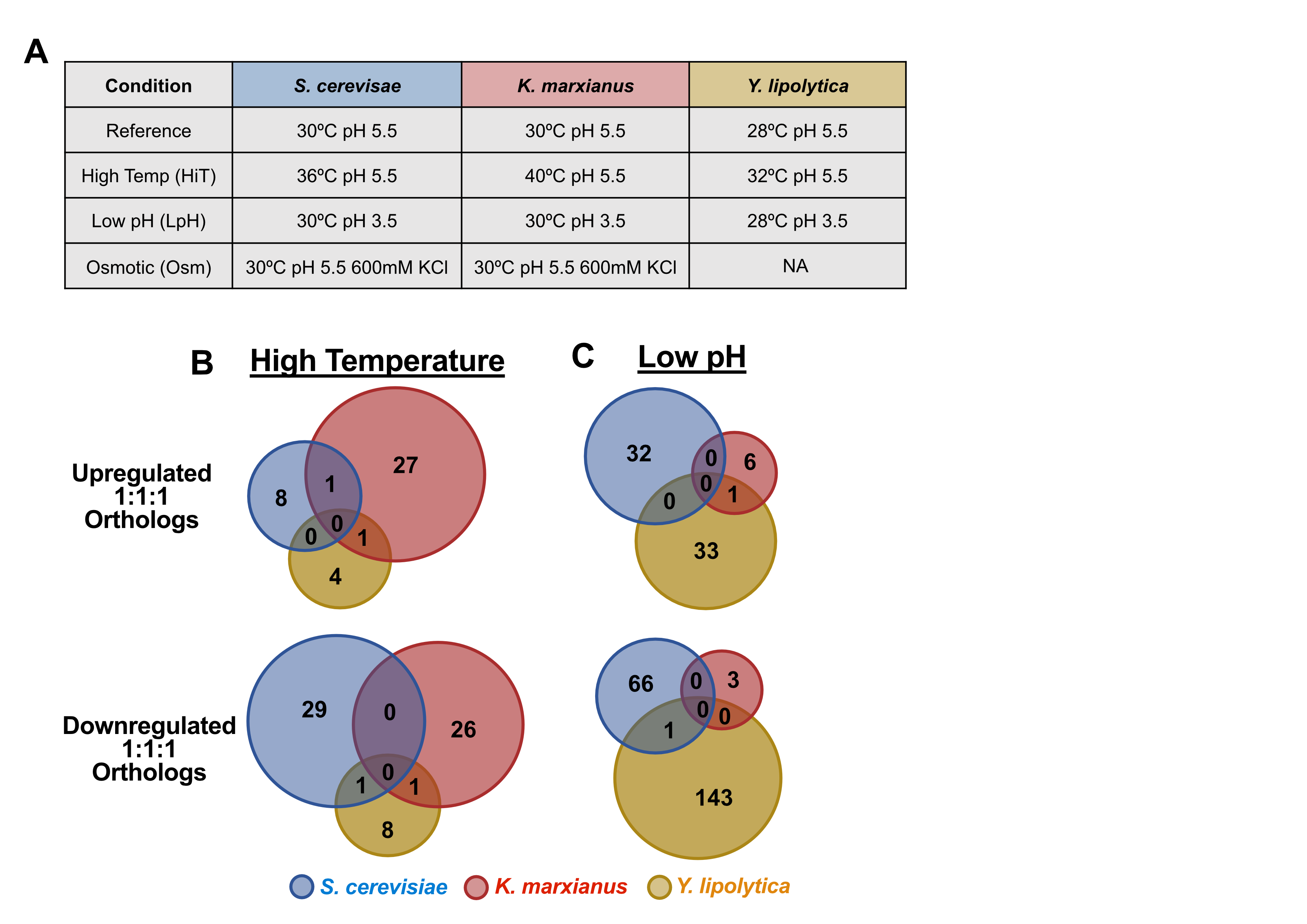

**Supplemental Figure 8) Single-Copy Core Orthologs Share Few Significant Gene Expression Changes in Response to Stress**

A. Chemostat conditions for each yeast species analyzed in this manuscript are shown. Differentially expressed orthologs are shown (log2FC>1, FDR<0.01), downregulated single-copy orthologous genes (bottom) and upregulated genes (top) for high temperature (B) and low pH (C). Overlapping regions indicate differential expression of an orthologous gene in each organism in the stress condition.

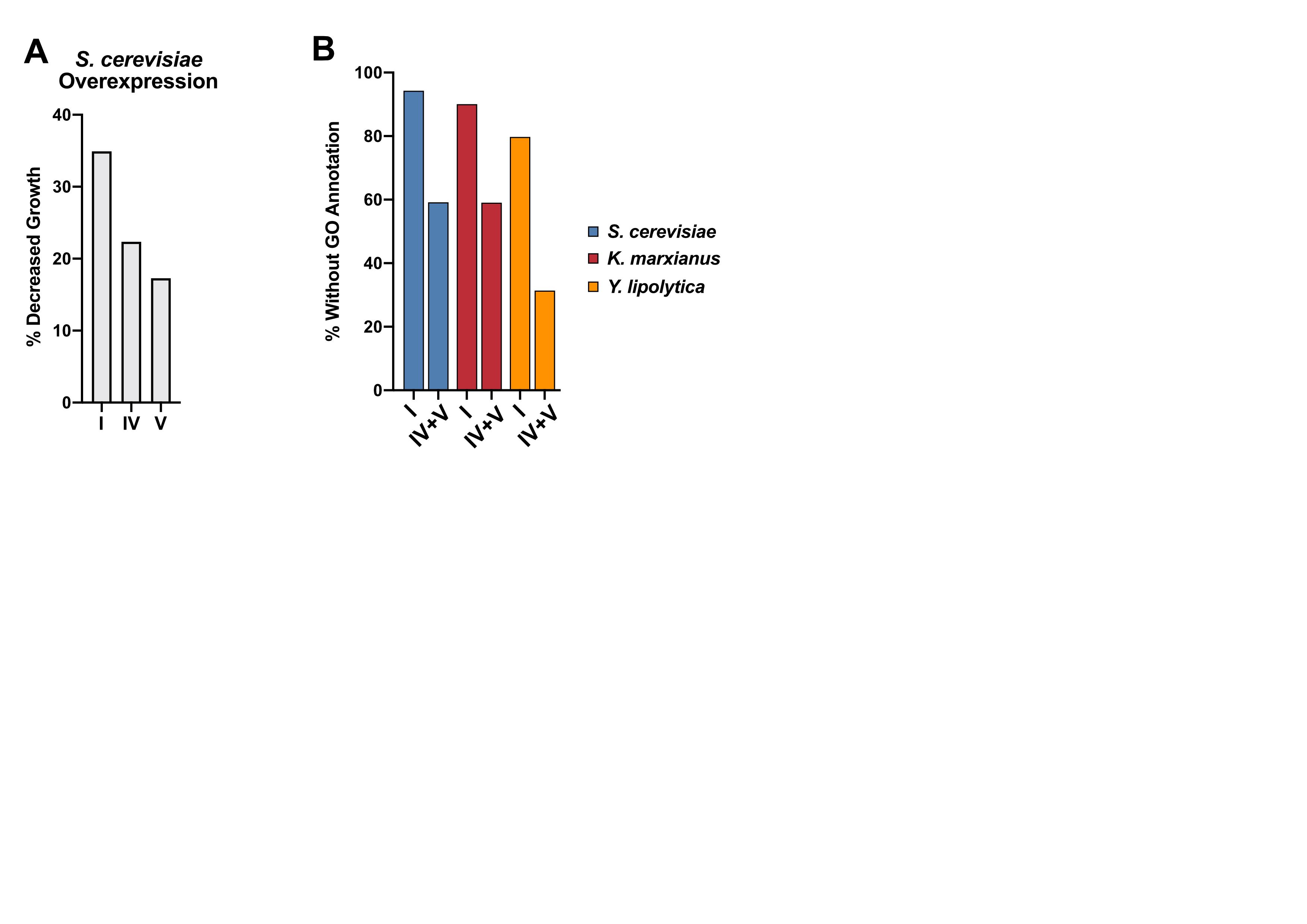

**Supplemental 9) Young Genes are Less Likely to Inhibit Growth Upon Overexpression and Often Lack Biological Process Information**

A. The percentage of genes exhibiting slower growth upon overexpression. Gene overexpression data was obtained from Vakirilis *et al.,* 2019^8^. B. The percentage of ancient (group I) and young (groups IV and V) genes with at least one biological process GO term. GO terms were acquired from Ensembl (*S. cerevisiae*) or from homology search via BLAST2GO^1^ (*K. marxianus* and *Y. lipolytica*).

**Supplementary Methods:**

**Gene Sorting Analysis Pipeline for Figures 2, 3, and Supplemental Figure 7**

Gene sorting analyses were based on ortholog inference which uses amino acid sequence similarity and chain length similarity to match proteins to ortholog groups from proteome versus proteome comparisons^3^. In this report, the protein orthogroups from each of these queries were matched to their corresponding genes for gene expression comparison. Queries were executed with the species in question (e.g. *S. cerevisiae*), and the sets of three species shown in Supplemental Figure 4B. Gene grouping was performed using an R-script that considers the raw outputs from OrthoFinder^3^, an example of raw data files can be sorted using the pipeline at: <https://github.com/SysBioChalmers/OrthOmics>. The gene grouping results for the genes measured in this study can be found in Supplemental Sheet II.

For *S. cerevisiae*, gene sorting first separated whole genome duplication genes/proteins identified in a previous work^4^ from the proteome. The timing of this event has been traced to a distinct evolutionary event that occurred ~93 million years ago^6^, thus, we used this information to isolate these as genes involved in a known evolutionary event. The remaining proteins were subject to an orthology search using OrthoFinder that compared the query species proteome to itself (e.g. *S. cerevisiae*) to identify single copy and duplicated proteins. This analysis identified 4,351 proteins as single copy and 115 proteins as non-WGD duplicates, which shared high homology and similar length to at least one other non-WGD protein from *S. cerevisiae*. These two groups were then subject to the gene sorting methods described below.

Single copy proteins were matched to groups based on the most distant ancestor that encoded an orthologous protein. For example, if a single copy protein from *S. cerevisiae* matched to any orthologous protein(s) from any ancestor in group I, then the protein was added to group I (and only group I). If a single copy protein did not match to an ortholog in group I, then the protein is screened for presence in any ancestor in group II (as shown in Supplemental Figure 4). This process continued until all single copy proteins were placed into a group with a matching ancestral gene or were not matched to any ancestors (species-specific groupV). This grouping paradigm is similar to phylostratigraphy^9^, and seeks to trace the relative timing of a gene origin event for single-copy genes. The same sorting paradigm was used to sort single copy genes from *K. marxianus* and *Y. lipolytica*.

For multi-copy proteins, sorting combined information about gene origin (above) with the added requirement copy number conservation. This forces co-migration of all duplicated family members of multi-copy protein families into a single group. Sorting of multi-copy protein families was done via a bottom-up approach, which considered whether the duplication was present in members of the genus first (Supplemental Figure 4D). An example is the duplicate protein family Oye2/Oye3, which were found to be duplicated in other members of the genus, clade, subphylum, and phylum, suggesting that this duplication likely occurred prior to the origin of the budding yeast subphylum. Oye2/Oye3 were therefore added to group I, as these are ancient genes that duplicated 400 million years ago or more. A counter example is the protein family Pma1/Pma2, which were found to be duplicated in members of the *Saccharomyces* genus but were not involved in the whole genome duplication and were not found to be duplicated amongst members of the Saccharomycetaceae clade. Although the ancestral gene for this family of plasma membrane ATPases is found broadly amongst yeasts, including filamentous fungi, the duplication event resulting in Pma1 and Pma2 is inferred to have occurred after the *Saccharomyces* genus emerged, and thus, these proteins were added to group IV.

**Proteomics Sample Collection and Preparation**

Samples for proteome analysis were collected as in de Groot *et al.,* 2007 with minor variations^10^. After 2x20mL aliquots collection for each chemostat, the samples were kept on ice and immediately centrifuged (12 000 g, 7 min at 0°C), washed once with ice-cold milliQ water, and stored as a pellet at -80°C.

Pellets were washed twice with 50 mM TRIS-HCl pH=7.8 and suspended in 3 mL of lysis buffer that contained 6 M urea (Sigma Aldrich, U5378), 2 M thiourea (Sigma Aldrich, T8656), 5 mM DTT (Sigma), 0.1 M TRIS-HCl pH=8 and 150 µL of prepared protease inhibitor cocktail (Sigma Aldrich, P8465). Cells were disrupted using a cell disruptor (Constant systems Ltd. One shot model) at 2.4 Kbars and cell debris was removed by centrifugation (15 min at 4000 g, 4°C). Soluble proteins were precipitated following a trichloroacetic acid (TCA)-acetone protocol^11^. Proteins were resuspended in lysis buffer and total protein concentrations were measured according to the 2-D Quant kit protocol (GE Healthcare Life Sciences, 80-6483-56). Aliquots of 80 µg of total protein extract suspended in 60 µL of lysis buffer were supplemented with RapiGestTM (Waters, 186001860) to a final concentration of 0.1% m/v. This protein extract was used for the proteome analysis.

The total protein extracts (80 µg) were alkylated with iodoacetamide (50 mM final concentration) in the dark for 45 min. Samples were first digested in-solution for 3 h at room temperature by adding PierceTM LysC-Protease (ThermoFisher Scientific, 90051) at a 1:50 (w/w) protein ratio. Then, following a 6-fold dilution with deionized water, a second overnight digestion was performed with 1:50 (w/w) sequencing-grade modified trypsin (Promega) at 37°C10.

To quench the digestion, the pH of the peptide mixtures was adjusted to 2 by adding 4 µL of trifluoroacetic acid (TFA) 30% v/v. The resulting peptide mixtures were pre-cleaned with a Strata-X column (Phenomenex, ref. 8B-S100-TAK). Columns were washed with 1.5 mL of washing buffer (containing 3% acetonitrile (ACN) and 0.06% glacial acetic acid). The peptide mixtures were charged into the columns, followed by three washing steps of 500 µL. Elution of peptides was achieved using 600 µL of elution buffer (40% ACN and 0.06% glacial acetic acid). The resulting samples were concentrated under vacuum to dryness and resuspended in 320 μL of loading buffer (0.08% TFA and 2% ACN) for analysis in a high-resolution mass spectrometer.

**LC-MS/MS analysis**

MS analyses were performed on a Dionex U3000 RSLC coupled to an Orbitrap Fusion™ Lumos™ Tribrid™ mass spectrometer (Thermo Fisher Scientific). Four μL containing 1 μg of digested peptides were injected and separated using a packed column claim PepMap®, 75 μm x 500 mm, C18, 3 μm, 100 Å, (Thermo Fisher Scientific). Buffer A consisted of 0.1% formic acid in 2% ACN and buffer B of 0.1 % formic acid in 80% ACN. The peptide separation analysis was achieved at 300 nL/min with a linear gradient from 1 to 35% buffer B for 160 min and 35% to 50% for 10 min. One run took 195 min including the regeneration step at 98 % buffer B. Ionization (1.6 kV ionization potential) and capillary transfer (275°C) were performed with a liquid junction and a capillary probe (SilicaTip™ Emitter, 10 μm, New Objective).

MS/MS analysis was performed in data dependent acquisition mode, with a top speed cycle of 3 s for the most intense double or multiple charged precursor ions. Ions in each MS scan over threshold 50,000 were selected for fragmentation (MS2). The mass spectrometer acquisition settings were set as follows. Full MS scan in Orbitrap (scan range [m/z] = 400–1600) with a resolution of 120,000 (AGC target = 5 x 105, max. injection time of 100 ms, data type = centroid). Analyzed charge states were set to 2-5 with a top speed cycle of 3s for the most intense double or multiple charged precursor ions. The dynamic exclusion within 10 ppm during 60 s and the intensity threshold was fixed at 5 x 104. And MS/MS using High Collision Dissociation (HCD) in the Orbitrap with resolution of 15,000 (30% collision energy, AGC target of 5.0 x 104 and max. injection time = 54 ms). Polysilaxolane ions m/z 445.12002, 519.13882 and 593.15761 were used for internal calibration.

**Protein identification**

Database searches were performed using the X!Tandem algorithm (version Alanine 2017.02.01; http://www.thegpm.org/TANDEM/) implemented in the open source search engine X!TandemPipeline version 3.4.4^12^. Enzymatic cleavage was declared as a trypsin digestion with one possible mis-cleavage. Carboxyamidomethylation of cysteine residues and oxidation of methionine residues were set to static and possible modifications, respectively. Precursor mass tolerance was 10 ppm and fragment mass tolerance was 0.02 Th. Identified proteins were filtered and grouped using X!TandemPipeline. Data filtering was achieved according to a peptide E-value < 0.05, protein log(E-value) < –2 and to a minimum of two identified peptides per protein. Using such filtering criteria, the peptides and proteins false discovery rates (FDR) were estimated between 0.02-0.3% and 0.04-0.11 %, respectively. MS data were deposited online on the public databases PROTICdb^13,14^available in the following URLs: http://moulon.inra.fr/protic/chassy_kluyveromyces, http://moulon.inra.fr/protic/chassy_yarrowia and http://moulon.inra.fr/protic/chassy_saccharomyces; as well as to the ProteomeXchange Consortium via the PRIDE^15^ partner repository with the dataset identifier PXD011426.

These methods were able to quantify 38.6% (2100 proteins), 32.9% (1684 proteins) and 27.5% (2186 proteins) of the theoretical proteome of *S. cerevisiae* CEN.PK113-7D, *K. marxianus* CBS 6556, and *Y. lipolytica* W29, respectively.

**Data Analysis Pipeline**

An integrated toolbox for analysis of RNAseq and proteomics data that incorporates GO terms annotation and gene orthology information (from OrthoFinder) has been generated in R programming language. Transcripts and proteins were considered as detected if they were measured in at least 2/3 of the total replicates for a given condition, an additional filter removes reads with relative standard deviation greater than 1 (RSD>1) across its replicates.

Relative proteomics measurements (XIC) were divided by the molecular weights of the proteins in order to take the overrepresentation of large proteins in the datasets.

Differential expression analysis of RNAseq data was carried out by using limma^16^ and edgeR^17^ R packages. Differential expression of genes was defined by a significance cutoff of absolute log2FC>1 for a stress condition compared to control, and an adjusted p-Value lower than 0.01, adjusting for multiple testing^18^.

The multi-omics data analysis toolbox has been generated using Git version control system, and it is available as a public repository at: https://github.com/SysBioChalmers/OrthOmics. Raw data files and other visualization tools and outputs can also be found in the repository.
